# Comparative population genetic structure of two ixodid tick species (*Ixodes ovatus* and *Haemaphysalis flava*) in Niigata Prefecture, Japan

**DOI:** 10.1101/862904

**Authors:** Maria Angenica F. Regilme, Megumi Sato, Tsutomu Tamura, Reiko Arai, Marcello Otake Sato, Sumire Ikeda, Maribet Gamboa, Michael T. Monaghan, Kozo Watanabe

## Abstract

Ixodid tick species such as *Ixodes ovatus* and *Haemaphysalis flava* are essential vectors of tick-borne diseases in Japan. In this study, we investigated the population genetic structures and gene flow of *I. ovatus* and *H. flava* as affected by the tick host mobility. We hypothesized that *I. ovatus* and *H. flava* may have differences in their genetic structure due to the low mobility of small rodent hosts of *I. ovatus* at the immature stage in contrast to the mediated dispersal of avian hosts for immature *H. flava.* We collected 307 adult *I. ovatus* and 220 adult *H. flava* from 29 and 17 locations across Niigata Prefecture, Japan. We investigated the genetic structure at two mitochondrial loci (*cox1*, 16S rRNA gene). For *I. ovatus*, pairwise *F*_ST_ and analysis of molecular variance (AMOVA) analyses of *cox1* sequences indicated significant genetic variation among populations. Both *cox1* and 16S rRNA markers showed non-significant genetic variation among locations for *H. flava*. The Bayesian tree and haplotype network of *cox1* marker for *I. ovatus* samples in Niigata Prefecture found 3 genetic groups wherein most haplotypes in group 2 were distributed in low altitudinal areas. When we added *cox1* sequences of *I. ovatus* from China to the phylogenetic analysis, three genetic groups (China 1, China 2, and Niigata and Hokkaido, Japan) were formed in the tree suggesting the potential for cryptic species in the genetic group in Japan. Our results support our hypothesis and suggest that the host preference of ticks at the immature stage may influence the genetic structure and gene flow of the ticks. This information is vital in understanding the tick-host interactions in the field to better understand the tick-borne disease transmission and in designing an effective tick control program.

## Introduction

Tick-borne diseases are a public health concern, and their control is often challenging because of the complex transmission chain of vertebrate hosts and ticks interacting in a changing environment (Dantas-Torres et al., 2012). Population genetic studies helped understand the dispersal patterns and potential pathogen transmission in ticks (Araya-Anchetta et al., 2015). Population genetic studies can elucidate the direction, distance, and gene flow patterns between tick populations (McCoy, 2008). For example, if a high gene flow is observed, there might be a greater chance of colonizing new areas or re-colonizing areas following successful vector control programs.

Due to the small size of ticks and their vulnerability to harsh environments during off-host, tick dispersal is complex and is linked to its host movement (Falco and Fish, 1991). Host mobility can affect the genetic patterns of tick populations, although its effects are not consistent. Several studies have reported low gene flow in ticks with less mobile hosts (e.g., smaller mammals) and a high gene flow in ticks with highly mobile hosts (Araya-Anchetta et al., 2015). For example, *Amblyomma americanum* and *Amblyomma triste* (Acari, Ixodidae) exhibited a high gene flow across various spatial scales (137,000 km^2^ to 2.78 million km^2^) due to their hosts' dispersal capabilities (large mammals and birds) (Guglielmone et al., 2013; Mixson et al., 2006; Trout et al., 2010). Lampo et al., (1998) observed low gene flow in *Amblyomma dissimile* populations due to its hosts' low mobility (small mammals, reptiles, and salamanders).

In Japan, tick-borne diseases are an increasing public health concern, affecting humans and animals (Yamaji et al., 2018). A total of 8 genera of ticks have been recorded in Japan, composed of 47 species: 43 ixodid tick species and four argasid species (Fujita et al., 2006). Out of these 47 species, 21 species parasitize humans (Okino et al., 2010). Among these 21 species, *Ixodes ovatus*, the primary vector of the causative agents of Lyme borreliosis (Miyamoto et al., 1993), and *Haemaphysalis flava,* a vector of the causative agents of severe fever with thrombocytopenia syndrome (SFTS) and Japanese spotted fever (JSF), was reported in Japan (Yamaji et al., 2018; Yu et al., 2014). Yamaguti et al., (1971) observed that the hosts of adult *I. ovatus* were mainly hares but could also be large mammals (e.g., cows and horses) and that the hosts of immature forms were small rodents. They also found that the host preferences of adult *H. flava* were cows, dogs, horses, wild boar, bear, and deer, while the immatures parasitized birds. Comparative population genetic studies on two essential ixodid ticks: *I. ovatus* and *H. flava* are not existing.

In addition to population genetic structure, genetic analysis may also reveal the phylogenetic pattern and the presence of cryptic species, where morphologically identified individuals might represent more than one species (Fegan et al., 2005). Previous studies on *Rhipicephalus appendiculatus* (Kanduma et al., 2016), *Ixodes holocyclus* (Song et al., 2011), and *I. ovatus* (Li et al., 2018) have revealed the presence of cryptic species based on the clustering of haplotypes in a phylogenetic tree. Based on these previous studies, morphological criteria for species differentiation alone are equivocal and that genetic analysis is essential.

Here, we studied the population genetic structure of *I*. *ovatus* and *H*. *flava* and also tested for the presence of cryptic species using mitochondrial DNA sequences of *cox1* and the 16S rRNA markers. We hypothesized that *I. ovatus* and *H. flava* would display contrasting population genetic structures despite the overlap sharing of large mammalian hosts at the ticks’ adult stage. The low mobility of *I. ovatus’* host, mainly hares at the adult tick stage and small mammals during the immature stage may contribute to the ticks’ high genetic divergence. On the other hand, the combined high mobility of large mammalian host at *H. flava* adult stage and avian mediated dispersal at the immature stage may be affecting the ticks’ homogenized population genetic structure.

## Material and methods

### Study site, collection, sampling, and identification

From April 2016 until November 2017, ticks were collected by the standard flagging methods (Ginsberg et al., 1989) in 29 sites (Additional File 1. Table S1) across Niigata Prefecture in Japan. Ticks were collected 2 to 14 times in 6 core sites among the 29 sites (Additional File 2. Table S2). Altitude per location ranged from 8 from 1402 m/a.s.l (mean=350). The geographic distance between sites ranged from 4.28 to 247.65 km (mean=77.36). Collected ticks were stored in microcentrifuge tubes with 70% ethanol at 4°C. We identified the sex, developmental stage, and species using a compound microscope and identification keys of Yamaguti et al., (1971).

### DNA extraction, PCR amplification, and sequencing

Genomic DNA (*I. ovatus* n=320*; H. flava* n=220) from each identified adult tick individual was extracted using the Isogenome DNA extraction kits (NIPPON GENE Co. Ltd., Tokyo, Japan) following the manufacturer's recommended protocol. The total number of analyzed samples (n=540) was chosen based on its species identity, while the other identified tick species were excluded from this study. Before DNA extraction, each tick was washed with alcohol and a PBS solution. DNA concentration and quality were checked using a NanoDrop™ 2000 Spectrophotometer (Thermo Scientific™). Two mitochondrial genes were amplified by polymerase chain reaction (PCR): Cytochrome Oxidase 1 (*cox1*) (658 base pairs) using the primer pairs LCO-1490 (5’-GGTCAACAAATCATAAAGATATTGG - 3’) and HCO1-2198 (5’-AAACTTCAGGGTGACCAAAAAATCA-3’) (Folmer et al., 1994) and the 16S rRNA gene (16S) (407 base pairs) with the following primer pairs 16S+1 (5’CTGCTCAATGAATATTTAAATTGC-3’) and 16S – 1 (5’ - CGGTCTAAACTCAGATCATGTAGG-3’) (Tian et al., 2011). All PCR amplifications of both *cox1* and 16S were performed with a final volume of 10 µl with 1 µl of genomic DNA. The PCR reaction for both markers was composed of the following: 10x Ex Taq buffer, 25 mM MgCl_2_, 2.5 mM dNTP, 10 μm of forward and reverse primers, and five units/μl of TaKaRa Ex Taq™ (Takara Bio Inc.). The *cox1* PCR amplification was as follows: an initial denaturation of 94°C for 2 min, denaturation of 94°C for 30 s, annealing of 38°C for 30 s, an extension of 72°C for 1 min for 30 cycles, and a final extension of 72°C for 10 min. In contrast, the 16S PCR amplification followed the protocol of (Tian et al., 2011) with some modifications, initial denaturation of 94°C for 3 min, denaturation of 94°C for 30 s, annealing of 50°C for 40 s, an extension of 72°C for 40 s for 30 cycles and final extension of 72°C for 5 min. PCR products were purified using the QIAquick 96 PCR Purification Kit (Qiagen) following the manufacturer’s instructions and sequenced by Eurofin Genomics, Inc., Tokyo, Japan.

### Sequence data analysis

We assembled forward and reverse reads for each individual using CodonCode Aligner version 1.2.4 software (https://www.codoncode.com/aligner/). No ambiguous bases were observed, and the low-quality bases were removed at the start and end of the reads. Multiple alignments of sequences were performed using the MAFFT alignment online program with default settings (https://mafft.cbrc.jp/alignment/server/). To ensure sequence quality and correct species identification, we checked the sequences’ similarity against reference sequences from GenBank using BLASTN (Basic Local Alignment Search Tool – Nucleotide, https://blast.ncbi.nlm.nih.gov/Blast.cgi?PROGRAM=blastn&PAGE_TYPE=BlastSearch&LINK_LOC=blasthome). We checked the quality of the final aligned sequences (*cox*1 = 658bp; 16S = 407bp) in Mesquite version 3.5 (Maddison et al., 2011) protein-coding genes were translated into amino acids to confirm the absence of stop codons.

### Population genetic analysis

We combined the nearby sites with a geographic distance range of 8.83 km to 79.81 (mean=44.00) with less than 8 individuals for the population genetic analysis because the accurate estimation of allele frequencies is difficult for small size population. A total of 8 populations (A to H) (Additional File 1 Table S1) were formed. Some populations were not included in the population genetic analysis because of the small sample size (< 8 individuals) and can’t be combined with any nearby sites.

We analyzed the sequences of both *H. flava* and *I. ovatus* per genetic marker separately using DNASp version 6.12.03 (Rozas et al., 2017) and Arlequin version 3.5.2.2 (Excoffier and Lischer, 2010) for the following parameters: number of haplotypes (nh), the average number of polymorphic sites (s), and average number of nucleotide differences (k). The haplotype diversity (h) and nucleotide diversity (π) were calculated in Arlequin version 3.5.2.2. The population genetic structure among and within populations in Niigata Prefecture was assessed by analysis of molecular variance (AMOVA) performed in Arlequin with 9999 permutations. The degree of molecular genetic differentiation between populations was assessed by calculating the pairwise *F*_ST_ values using Arlequin.

To determine if the genetic differentiation was influenced by geographical distance or altitudinal differences among populations, we performed Mantel Test in GenAlEx version 6.51b2 (Peakall and Smouse, 2006). Two tests per species and marker were conducted. First, we compared pairwise genetic (pairwise *F*_ST_ values) and geographical distances (km), and second, we analyzed the genetic distance (*F*_ST_ values) with altitudinal differences (m/a.s.l.) calculated from GenAlex version 6.51b2. The geographic distances were obtained from the geographic midpoint of the populations using the GPS coordinates (latitude and longitude) of each site recorded during the sampling. The altitudinal differences are the calculated mean value per population. All Mantel tests were assessed using 9999 permutations for the significance of the correlation.

The genetic relationships among the populations were calculated by the unweighted pair group method with the arithmetic mean (UPGMA) cluster method using the APE package (Paradis and Schliep, 2018) and R program (R Development Core team., 2016). To create a dendrogram, we used the genetic distance matrix (pairwise *F*_ST_ values) generated from GenAlEx.

### Haplotype network and Phylogenetic analysis

To evaluate the haplotypes’ relationship, we constructed a haplotype network on the PopART program version 1.7 per marker (*cox1* and 16S) and species (*I. ovatus* and *H. flava* (http://popart.otago.ac.nz/index.shtml) using the median-joining (MJ) network algorithm (Bandelth et al., 1999).

We performed a Bayesian phylogenetic analysis using BEAST v1.10.4 (Drummond and Rambaut, 2007) to determine the phylogenetic structure of *I. ovatus* haplotypes within Niigata Prefecture using the *cox1* marker. Additional sequences from China were not included in the Bayesian analysis. We used the HKY substitution model with the estimated base frequencies. The strict clock model was employed and the speciation yule process as the tree prior. A maximum clade credibility tree was acquired using the TreeAnnotator v1.10.4 from the many trees obtained from BEAUti v1.10.4 with 10% of trees used during burn-in. The maximum clade credibility tree was viewed using FigTree v1.4.4.

We constructed maximum likelihood (ML) gene trees for *cox1* and 16S haplotype sequences of *I. ovatus* and *H. flava* using PhyML version 3.1 (Guindon and Gascuel, 2003) default settings. We applied HKY and GTR nucleotide substitution models for *cox1* and 16S, respectively, as suggested by jModelTest version 2 (Darriba et al., 2012). Additional sequences from China (MH208506, MH208512, MH208514, MH208522, MH208515-19, MH208524, MH208531, MH208574, MH208577, MH208579, MH208681-87, MH208689-93, MH208706, KU664519 (Li et al., 2018), Japan (Hokkaido AB231670, U95900; Yamanashi AB819241, AB819243 and Aomori AB819244) (Mitani et al., 2007; Norris et al., 1999; Takano et al., 2014) were included to know the presence of cryptic species. We used *Ixodes canisuga* as an outgroup because it is closely related to *I. ovatus* and *H. flava* (KY962023 and KY962074; Hornok et al., 2015).

## Results

A total of 2,374 individual ticks was collected. Adult and immature *Ixodes nipponensis*, *Dermacantor taiwanensis*, *Ixodes persulcatus*, and *Ixodes monospinus* were also identified and used for another research study. In this study, adult *I. ovatus* and *H. flava* were only analyzed because these two species were the most abundant. Larval ticks were used in another study. The number of *I. ovatus* ranged from 1 to 36 adult individuals per site and were more successfully sequenced for *cox1* (307/320; 95.9%) than for 16S (284/320; 88.8%). The analyzed *H. flava* ranged from 1 to 77 adult individuals per site and were also more successfully sequenced for *cox1* (220/223; 98.7%) than for 16S (172/223; 77.1%). On the other hand, for the population genetic analysis, each population (i.e., combined nearby sites) had the following tick individual range and mean per markers: *cox1 I. ovatus* (28 to 62; mean=43), 16S *I. ovatus* (24 to 66; mean=40), *cox1 H. flava* (8 to 81; mean=43) and 16S *H.flava* (8 to 76; mean=34). There were 60 and 63 *cox1* haplotypes and 24 and 40 16S haplotypes in *I. ovatus* and *H. flava,* respectively (Table 1). Haplotype diversity (h) ranged from 0.442 to 0.852, and nucleotide diversity (π) ranged from 0.001 to 0.006 for both markers and species.

**Table 1.** Summary of *cox1* and 16S haplotype, variability, genetic diversity of adult *I. ovatus* and adult *H. flava* populations in Niigata Prefecture, Japan

| Marker | Species | n | nh | s | k | h | $\pi$ |
| --- | --- | --- | --- | --- | --- | --- | --- |
| *cox1* | *I. ovatus* | 307 | 60 | 65 | 2.728 | 0.852 | 0.004 |
|  | *H. flava* | 220 | 63 | 60 | 1.472 | 0.789 | 0.002 |
| 16S | *I. ovatus* | 284 | 24 | 22 | 0.699 | 0.442 | 0.001 |
|  | *H. flava* | 172 | 40 | 49 | 2.447 | 0.835 | 0.006 |

Abbreviations: n sample size; nh number of haplotypes; s number of polymorphic sites; k mean number of nucleotide differences; h haplotype diversity; $\pi$ nucleotide diversity

AMOVA results revealed a high among-population divergence (61.99%) in *I. ovatus cox1* sequences, whereas both *cox1* and 16S markers of *H. flava* indicated no significant genetic variation among populations (Table 2). Significant global *F*_ST_ values (P<0.01) were observed for *I. ovatus* with significant values 0.3801 in *cox1* and 0.0378 in 16S markers. Pairwise *F*_ST_ values (0.0963 to 0.6808) of *cox1* marker for *I. ovatus* showed significant genetic differences between most pairs of populations such as between populations B and D and populations F and D (Additional File 3 Table S3). We also observed significant genetic differences in 16S *I. ovatus* sequences ranged from 0.0514 to 0.0949 (Additional File 4 Table S4). Pairwise *F*_ST_ values from the *cox1* marker of *H. flava* showed no significant genetic differences except in few population pairs such as A and C, C and E and D and E (Additional File 5 Table S5). No significant genetic differentiation was also observed in 16S marker of *H. flava* except between A and C (Additional File 6 Table S6).

**Table 2.**
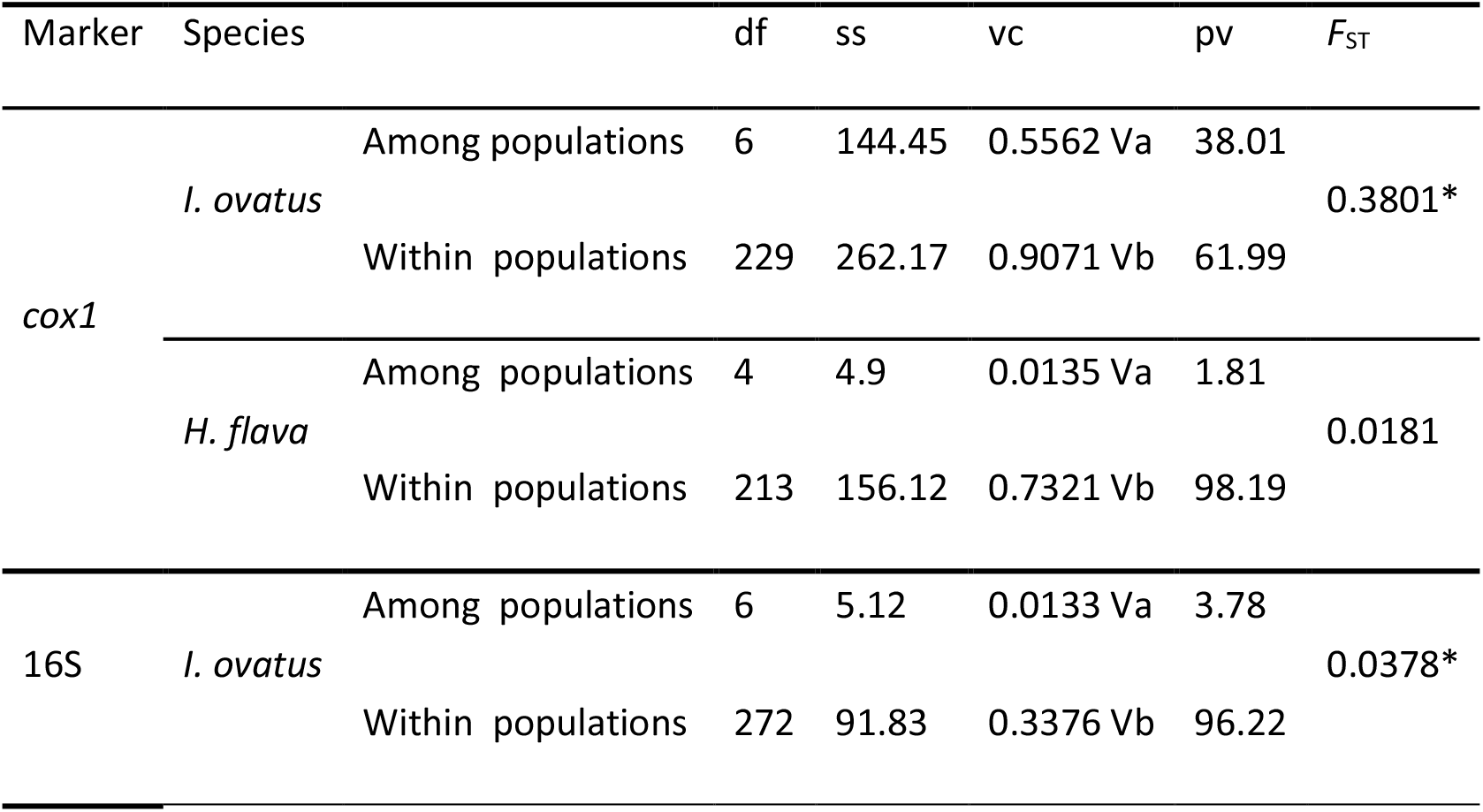
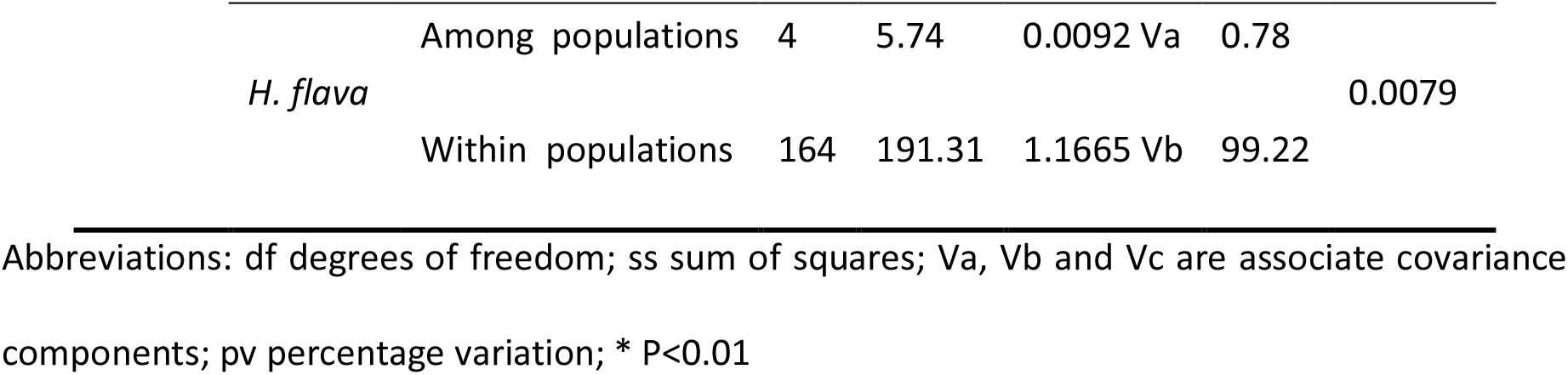
Analysis of molecular variance (AMOVA) using *cox1* and 16S of adult *I. ovatus* and adult *H. flava* populations

The Mantel test of *I. ovatus* showed no significant isolation by geographic distance in both markers: (*cox1 r* = 0.108, *p* = 0.269; 16S *r* = 0.518, *p* = 0.065) (Additional File 7 Figure S1, a and b) and no evidence for isolation by altitudinal difference (*cox1* r = −0.066, *p*=0.225; 1S r = −0.023, p = 0.577). Both Mantel tests were also not significant for *H. flava* (distance: *cox1* r= 0.444, p= 0.130; 16S r = 0.355, p= 0.189; altitude: r = 0.092, p = 0.30; 16S r = 0.217, p= 0.06) (Additional File 7 Figure S1, c and d).

The UPGMA cluster dendrogram constructed from the pairwise *F*_ST_ values of *I. ovatus* for the *cox1* marker (Figure 2) revealed two genetic clusters among the 7 populations, where cluster 2 included populations from the more western sites. In contrast, cluster 1 (A, B, and F) mostly had populations in the northern and southern parts of Niigata Prefecture (Figure 3) and are distributed in mountainous areas with high elevation. We observed no evident genetic clustering on the dendrogram of *I. ovatus* for 16S marker and those of *H. flava* for *cox1* and 16S makers, respectively (Additional File 8 Figure S2; Additional File 9 Figure S3; Additional File 10 Figure S4).

**Figure 1.**
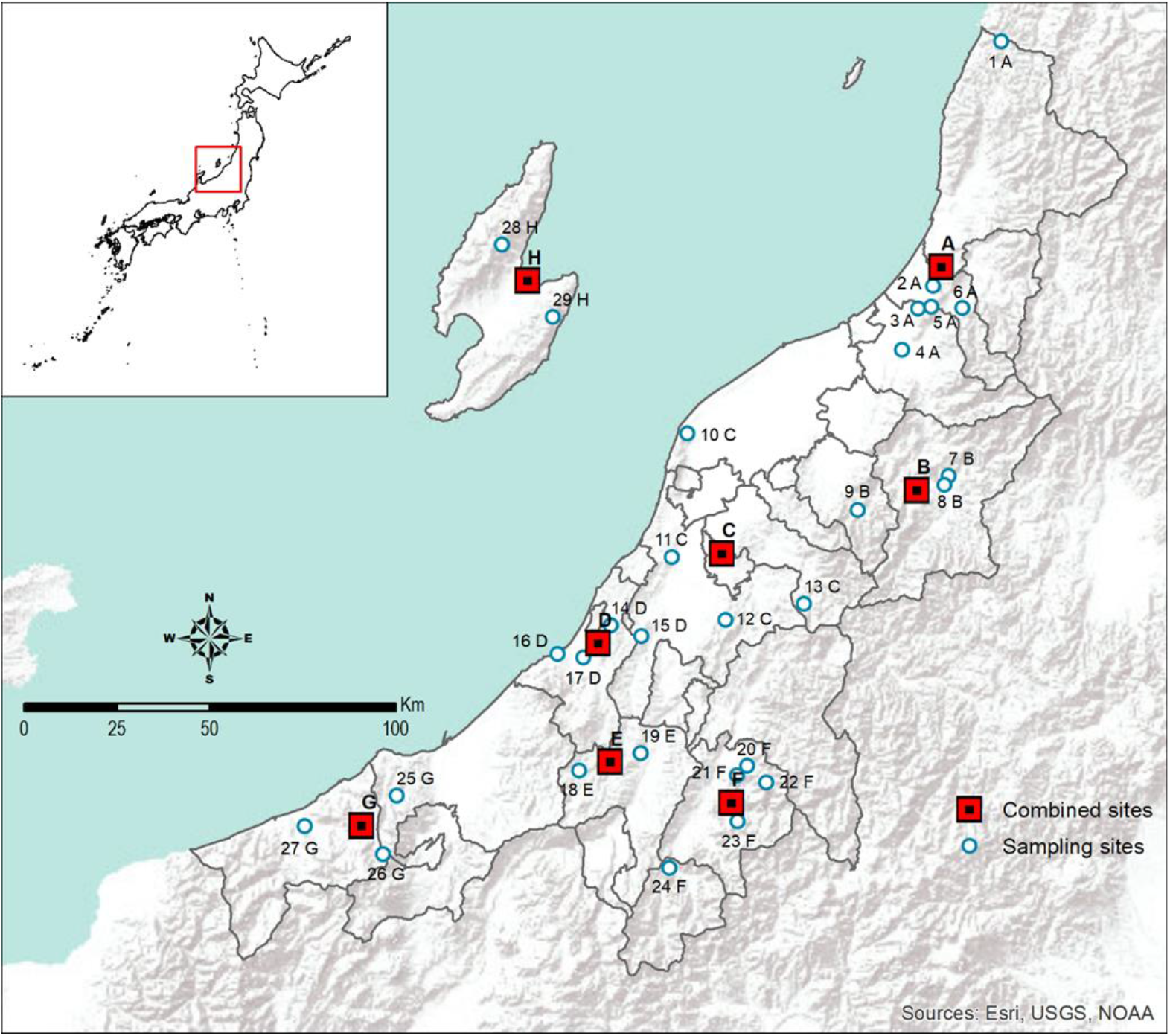
Map of the 29 sampling sites used for this study. The populations labeled A to H are composed of the nearby sites labeled 1 to 29 used for the population genetic analysis.

**Figure 2.**
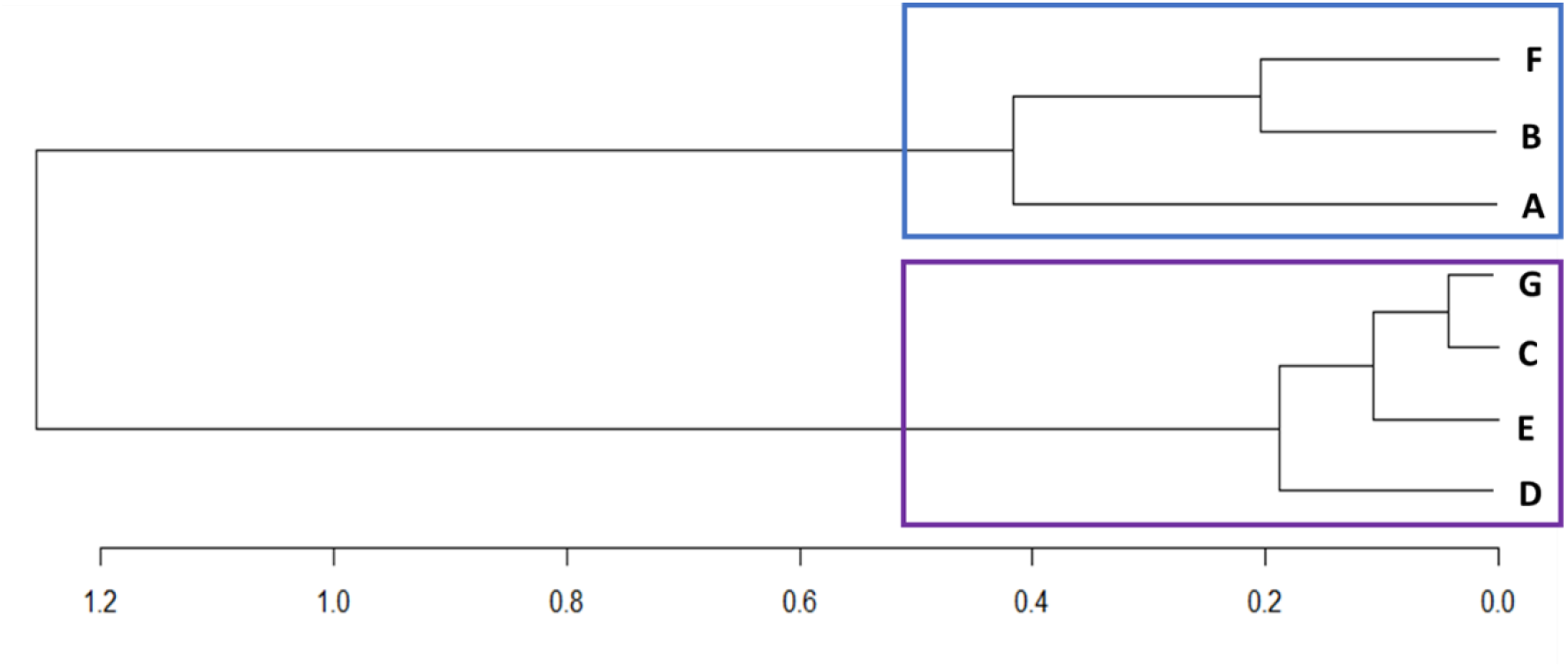
An unweighted pair group method with the arithmetic mean (UPGMA) dendrogram of adult *I. ovatus* based on the pairwise genetic distance (*F*_ST_) of *cox1* among the 7 populations across Niigata Prefecture, Japan.

**Figure 3.**
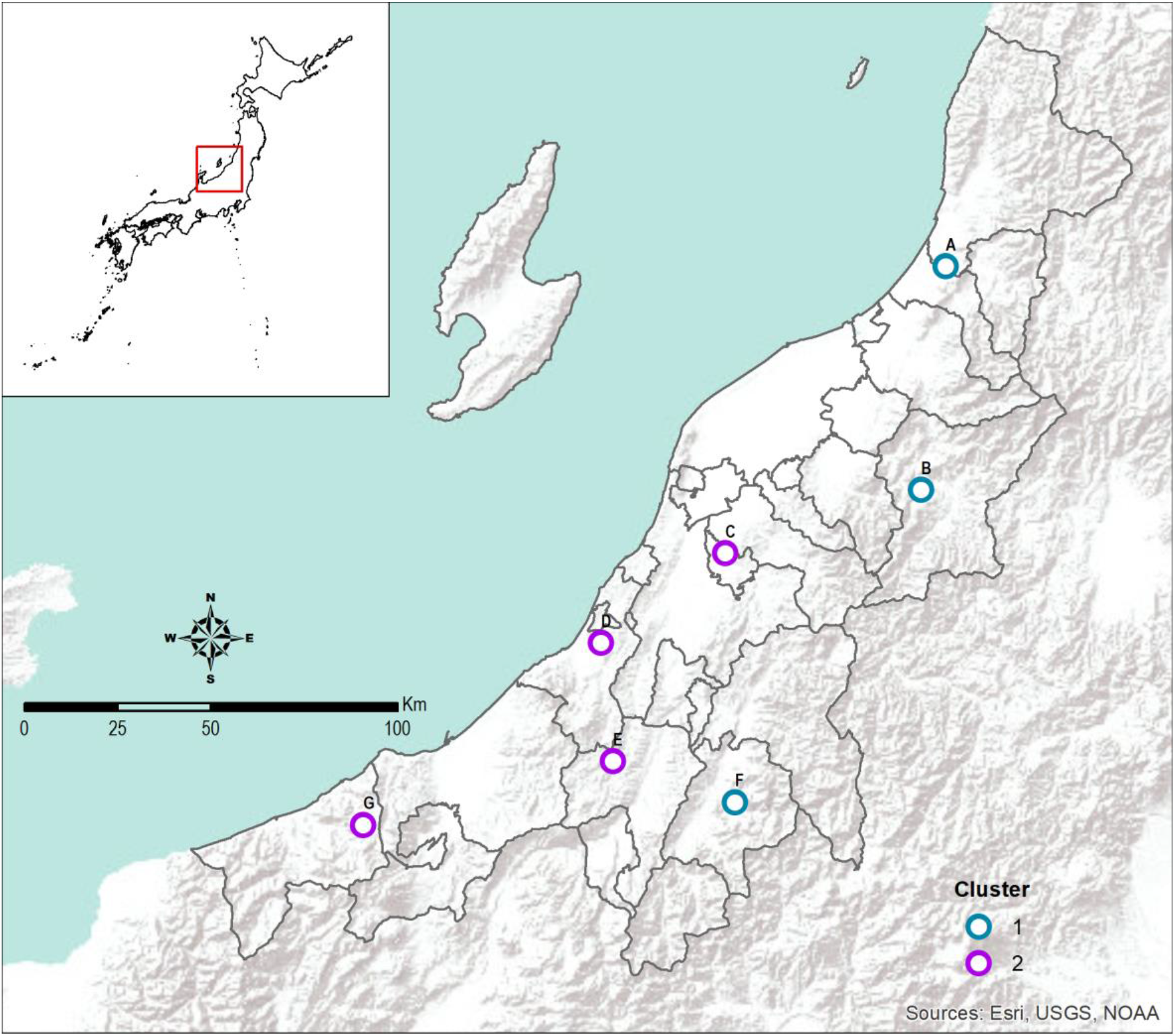
The distribution of the two genetic clusters as observed in the UPGMA cluster dendrogram (Figure 2) of *I. ovatus cox1* sequences from Niigata Prefecture.

The haplotype network with the reference sequences from China and the Bayesian tree of the *I. ovatus cox1* haplotypes showed similar patterns (Figure 5 and 6) of 4 genetic groups. The haplotype network of the *cox1* marker of *I. ovatus* also displayed the two distinct haplotypes (Hap 60 and Hap 59) (Figure 6; Additional File 11 Figure S5). These two haplotypes were found from sampling site 6 (Pop A) and sampling site 26 (Pop 6). The Bayesian tree of *I. ovatus cox1* haplotype sequences with reference sequences from China also displayed the 4 genetic groups (Additional File 12 Figure S6). The haplotype network of *I. ovatus* from 16S marker (Additional File 13 Figure S7), *H. flava* in both cox1 (Additional File 14 Figure S8) and 16S markers (Additional File 15 Figure S9) did not display any distinct haplotypes and genetic groups.

A putative *I. ovatus* species complex was identified in the phylogenetic tree based on three distinct haplotype groups found with both *cox1* and 16S rRNA markers; 2 groups in China and one in Japan. The *cox1* phylogenetic tree of 60 *I. ovatus* haplotypes indicated three groups: group 1 of the published sequences from western China, one large group 2 containing the 58 haplotypes and two divergent haplotypes (Hap 60 and Hap 59), and group 3 from west China (Figure 4). The published haplotype from Hokkaido, Northern Japan (Mitani et al., 2007) occurred within our large group, and the published sequences from western China were in a different group.

**Figure 4.**
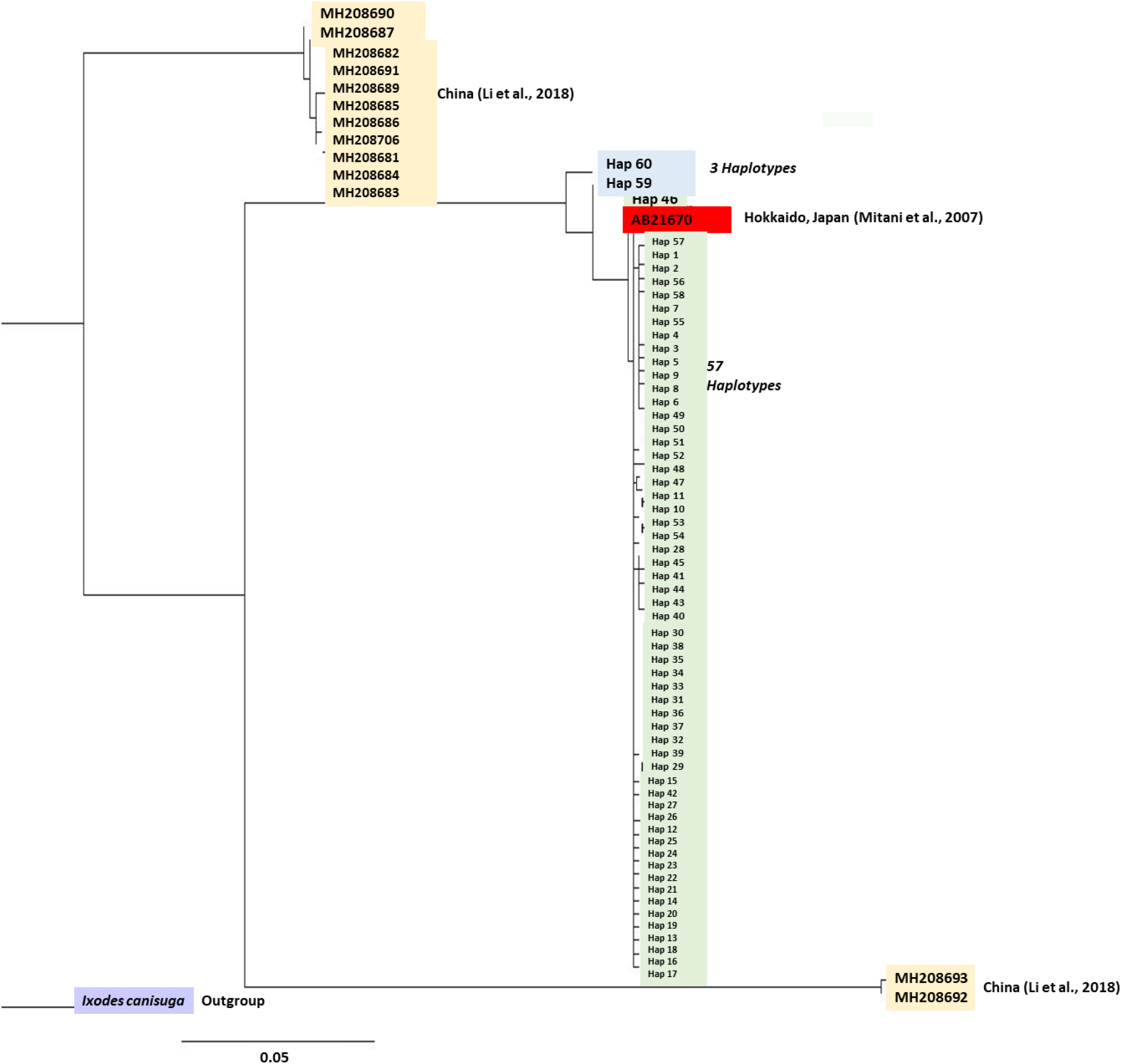
Maximum likelihood gene tree of *cox1* haplotype sequences of *I. ovatus* with sequences from China (MH208681-MH208687, MH208689-MH2086393, MH208706), Japan (AB231670), and an outgroup of *I. canisuga.* *Yellow* reference sequences from China (Li et al., 2018); *red* reference sequence from Hokkaido, Japan (Mitani et al., 2007); *blue I. ovatus* divergent *cox1* haplotypes; *green I. ovatus cox1* remaining haplotypes; *purple I. canisuga* outgroup

**Figure 5.**
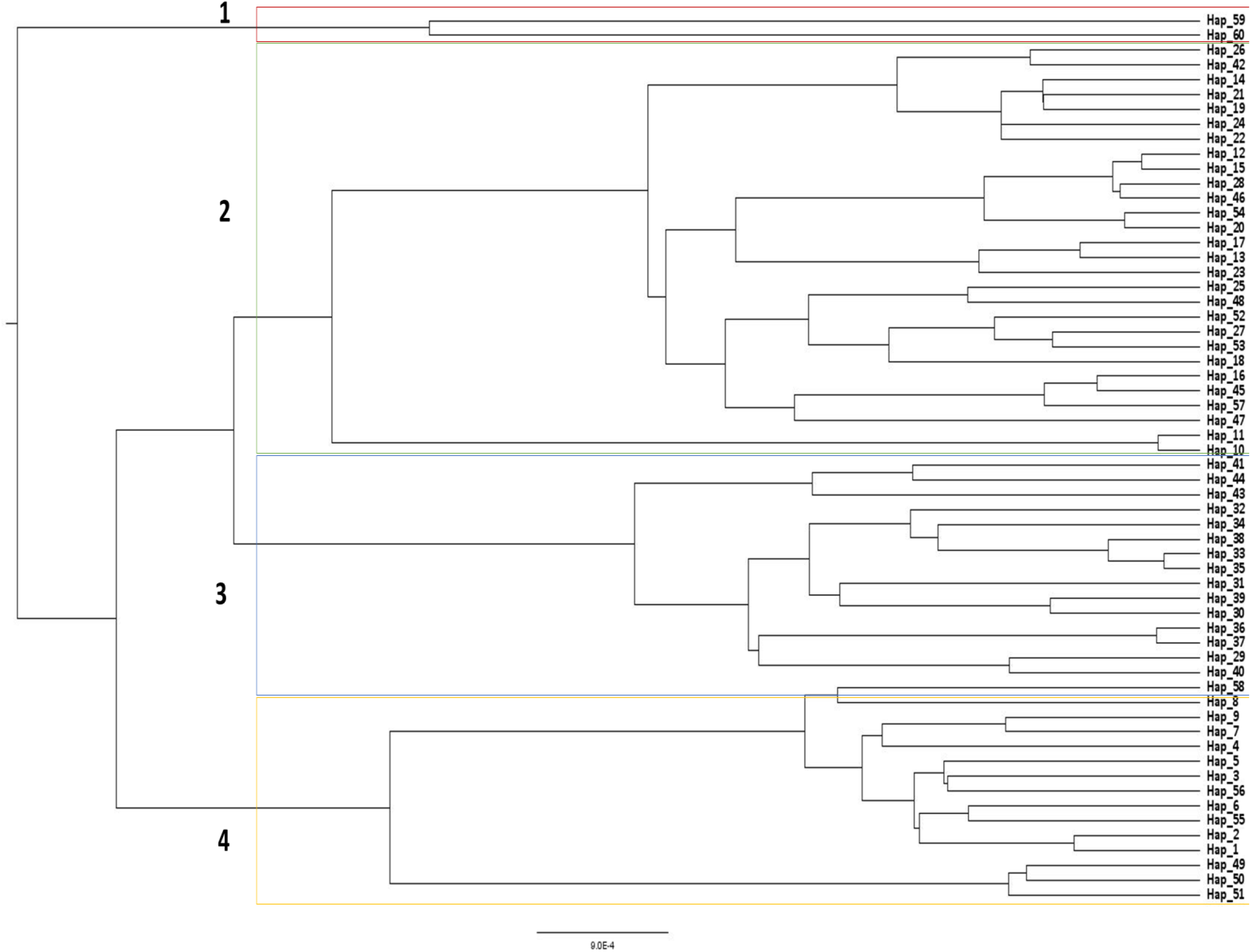
Phylogenetic tree from Bayesian analysis of the *cox1 I. ovatus* haplotype sequences

**Figure 6.**
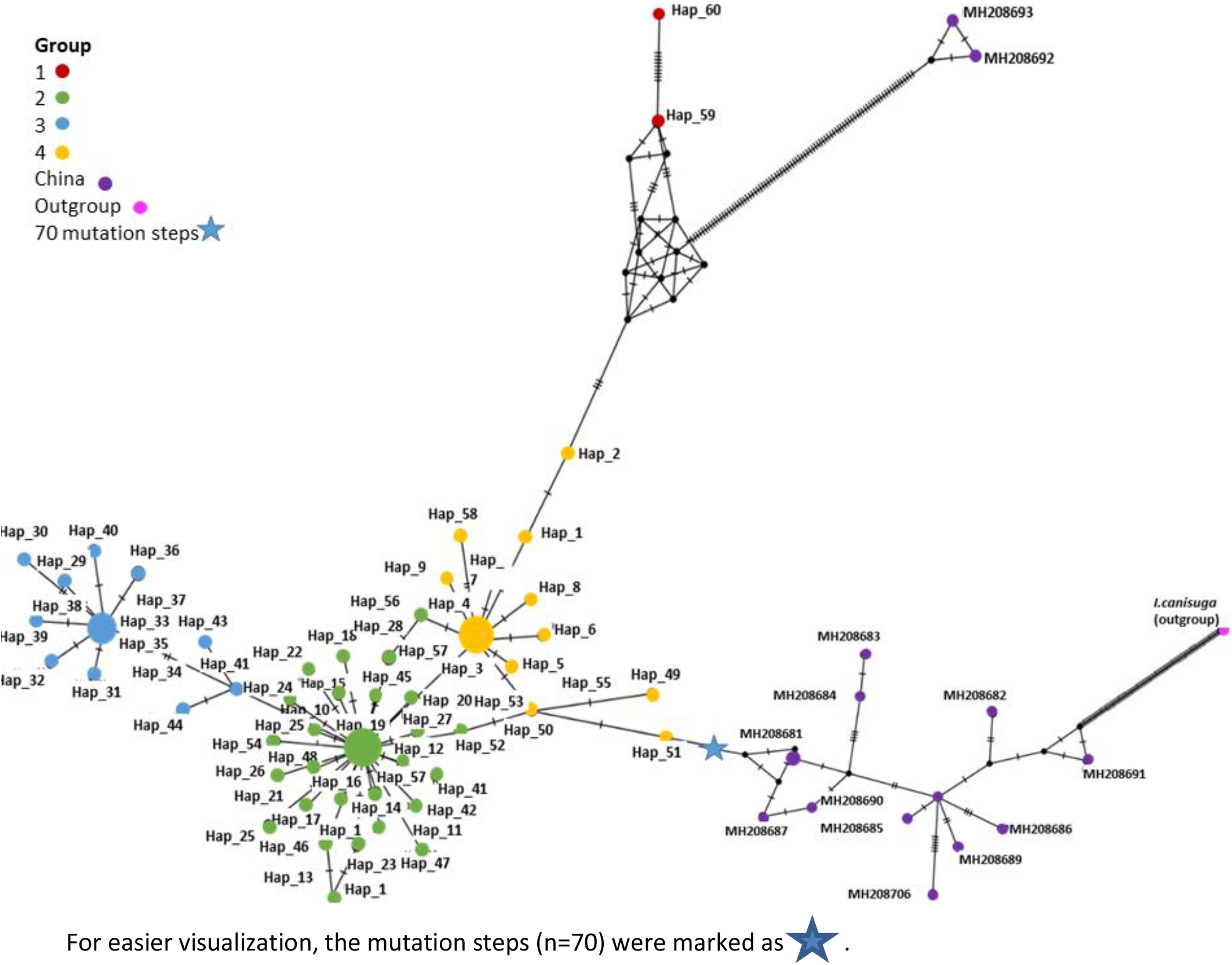
Median-joining haplotype network of *I. ovatus* haplotypes with reference sequences from China using the *cox1* marker. Each circle represents a unique haplotype and the lines correspond to mutations.

The *I. ovatus* haplotypes in the 16S phylogenetic tree (Additional File 16 Figure S10) were grouped with the published haplotypes from Japan (Yamanashi and Aomori Prefectures; Takano et al., 2014) and Hokkaido (Norris et al., 1999). The *cox1* haplotype sequences of *H. flava* phylogenetic tree (Additional File 17 Figure S11) were similar to reference sequences from China KY021800 – KY021807, KY021810 – KY021819 and KY003181 (Li, Z., Cheng, T. and Liu, G.; Unpublished results from NCBI); JQ625688 – JQ625689, JF758632 and JQ737097 (Lu et al. 2013) and JG737097 (Gou, H., Guan, G., Yin, H. and Luo, J.; unpublished results from NCBI). The *H. flava* 16S haplotype sequences (Additional File 18 Figure S12) showed high similarities with reference sequences from Japan (Kagoshima, Fukui Aomori, Fukui, Yamanashi, Kagawa, and Ehime Prefectures (Takano et al., 2014) and China KC844858 –KC844867 (Cheng et al., 2013); KX450280 –KX450282 (Zhang, Y., Cui, Y., Peng, Y., Yan, Y., Wang, X. and Ning, C. Liu, Q., Zhang, Y. and Zhu, D.; unpublished results from NCBI); MG696720 (Zheng, W., Chen, S. and Chen, H.; unpublished results from NCBI) and KP324926 (Liu, Q., Zhang, Y. and Zhu, D.; unpublished results from NCBI) sequences from GenBank.

## Discussion

### Contrasting population genetic structures between *I. ovatus* and *H. flava*

Our results supported our hypothesis that *I. ovatus* may display high genetic divergence because of its low host mobility in contrast to the homogenized population genetic structure in *H. flava* due to the combined large mammalian hosts at the ticks’ adult stage and avian mediated dispersal during the ticks’ immature stage. The significant *F*_ST_ estimates in *I. ovatus cox1* (0.3801) and 16S (0.0378) revealed population differentiation as supported by the high among populations percentage variation (61.99) in the AMOVA analysis. The high genetic divergence in *I. ovatus* can be due to the low mobility of its hosts, for example during the ticks’ adult stage wherein they utilized mostly hares while small rodents at the immature stage of *I. ovatus.* The host preference of *I. ovatus* may also contribute to the observed 4 genetic groups in the Bayesian tree and the haplotype network analysis. The haplotypes from groups 2 and 3 were mostly from low altitudinal areas as compared to the haplotypes from group 1 with ticks from the high altitudinal area such as in mountain areas. We assume that the large mammalian hosts such as cows and horses of *I. ovatus* have enabled the ticks to reach high mountain ranges from group 1. The distribution of the type of hosts may also affect altitude affecting also the formation of groups 1, 2, and 3 in the *cox1 I.ovatus* dendrogram.

On the other hand, the homogenized population genetic structure observed in *H. flava* might be because of the combined high host mobility of large mammals at the ticks’ adult stage and the avian mediated dispersal at the immature stage. Twenty-eight species of birds were previously reported as hosts of immature *H. flava* from Japan, mostly from the order Passeriformes (Yamauchi and Takeno, 2000). Large mammals and birds may have expansive habitats ranges that may allow high gene flow of *H. flava* between the locations in Niigata, as previously observed in *Amblyomma americanum* populations (Mixson et al., 2006; Reichard et al., 2005; Trout et al., 2010) and *I. ricinus* (Casati et al.,2008).

Several alternative factors such as tick behavior, biology, and ecology can also affect tick’s dispersal patterns. A previous study on two tick species: *Hyalomma rufipes* and *Amblyomma hebraeum,* also displayed contrasting genetic patterns despite the overlap sharing of highly mobile hosts such as birds (Cangi et al., 2013). This might be due to the species specific capacity of the immature stages of ticks to survive even after host detachment in various habitat conditions (Cumming, 1999; Estrada-Peña, 2015; Needham et al., 1986). Population genetic structure can also be influenced by assortative mating (e.g., *I. ricinus*) wherein genetically similar individuals tend to mate rather than random mating resulting in decreased genetic divergence (Kempf et al., 2009). Our study does not have supporting data to test these alternative factors; thus, it is suggested in future studies to analyze further these factors that could play a role in the tick’s dispersal movement.

16S marker showed significant genetic differentiation of *I. ovatus* (pairwise *F*_ST_ =0.0514 to 0.0949) in few pairs of sampling locations as compared to the *cox1* marker, which may be due to lower variability of the marker thus providing inadequate genetic variation for population genetic analysis, as previously observed in *Amblyomma ovale* (Bitencourth et al., 2019) and *Rhipicephalus microplus* (Burger et al., 2014; Low et al., 2015). The low variability of 16S marker in *I*. *ovatus* may not be sufficient to infer an intraspecific relationship and is further supported by its low (*nd* = 0.001) nucleotide diversity as compared to *cox1* marker (*nd* = 0.004). Thus, it may also explain the grouping observed in the *cox1* phylogenetic tree of *I. ovatus* was absent in the 16S tree. Despite these, we used the 16S marker because this gene mutates at a rate that is informative for inter-species level phylogenetic analysis (Araya-Anchetta et al., 2015). The negative pairwise *F*_ST_ values observed in *H. flava* do not have any biological meaning. These results from the unbiased estimator allowing a negative *F*_ST_ when the sample size is small. (Bitencourth et al., 2016; Vial et al., 2006; Weir and Cockerham, 1984).

### Species complex formation in *I. ovatus cox1* sequences

Our *cox1 I. ovatus* gene tree showed Japanese individuals to form a distinct group from haplotypes from southern China (Li et al., 2018). Despite the low gene flow of *I. ovatus* in *cox1* marker, we found that Niigata’s haplotypes were near related to the published sequence from Hokkaido in northern Japan. The result indicates that these ticks originated from a diverse set of geographical locations in Japan, which might be transported by its hosts or are undergoing recent population expansion from northern Japan (Hokkaido) to central Japan (Niigata) *vice versa*. We found three groups (China 1, Japan, and China 2) and two slightly divergent *cox1* haplotypes (Hap 60 and 59) in the Japan group of *I. ovatus* in Niigata. Considering two or more cryptic species can be concealed in one morphologically described species (Bitencourth et al., 2016), the occurrence of the three groups and the divergent haplotypes suggests that *I. ovatus* may be a species complex. It can be inferred that China 1 and China 2 might have a longer evolutionary time as compared to Japan. The co-existence of a species in the same geographic area may explain the occurrence of the species complex. One of the limitations of this study is that both Chinese and Japanese individuals were collected in a limited geographical area. We suggest that future studies sample and sequence more individuals from other locations. Extensive geographical sampling can explain better the taxonomy and the genetic structure of species as suggested in the study of Liu et al., (2013) on *Rhipicephalus sanguineus*. Previous studies have also observed species complexes in *Ixodes* and *Rhipicephalus* (Burger et al., 2014; Li et al., 2018; Song et al., 2011; Xu et al., 2003), suggesting that morphological criteria for species differentiation alone are equivocal and that genetic analysis is essential. Future studies must tackle an integrative approach of the tick species concept that includes the morphology, genetics, biology, and ecological trails of ticks (Dantas-Torres et al., 2012).

## Conclusions

In summary, our findings revealed the contrasting patterns of the genetic structure of *I. ovatus* and *H. flava* from Niigata Prefecture, Japan. The high genetic divergence in *I. ovatus* might be influenced by the restricted movement of its small mammalian hosts during the ticks’ immature stage and hares at the adult stage. On the other hand, the homogenized genetic structure in *H. flava* might be due to the avian mediated movement during the ticks’ immature stage coupled with the expansive movement of the large mammalian host at the ticks’ adult stage. Our results suggest that the host preference of ticks during its immature stage may strongly influence the population genetic structure of ticks due to its higher ability to survive in an environment. Understanding the population genetic structure of ticks such as *I. ovatus* and *H. flava* can give information about the distribution and control of tick-borne diseases. Even though *I. ovatus* populations were genetically structured within Niigata, a published haplotype from Hokkaido was also found, indicating that widespread dispersal is possible. The occurrence of three groups and the divergent *cox1* haplotypes in *I. ovatus* emphasizes the need for additional methods in determining the existence of a species complex in *I. ovatus* populations in Japan.

## List of Abbreviations

*H. flava*: *Haemaphysalis flava*
*I. ovatus*: *Ixodes ovatus*
AMOVA: analysis of molecular variance
UPGMA: unweighted pair group method with arithmetic mean
ML: maximum likelihood
*F*_ST_: fixation index
PBS: phosphate-buffered saline

## Declarations

### Ethics approval and consent to participate

Not applicable

### Consent for publication

Not applicable

### Availability of data and materials

All data generated or analyzed during the study are included in this published article and its additional supplementary files. All the newly generated sequences are available in the GenBank database under the accession numbers MW063669-MW064124 and MW065821 - MW066347.

### Authors’ contributions

MAFR, MS, and KW conceptualized and designed the experiment. MS, TT, RA, SI, and MOS designed the sampling collection, collected and identified the tick samples. MAFR and MG conducted the molecular analyses. MAFR, MM, and KW performed the data analysis. MAFR and KW wrote the manuscript. All authors read and approve the manuscript.

## Acknowledgments

The authors would like to thank the Niigata Prefectural Office for their valuable help during the tick sampling collection. We are also thankful to Masaya Doi, Kohki Tanaka, and Mizuki Ueda for their technical assistance in the molecular analyses. Thank you to Dr. Thaddeus M. Carvajal and Dr. Joeselle M. Serrana for their useful suggestions in the population genetic analyses in this study and Micanaldo Francisco for technical assistance in constructing this study’s sampling map.

## Additional Files

**Additional File 1.xls Table S1**. Summary of adult *Haemaphysalis flava* and *Ixodes ovatus* collected from the different locations of Niigata Prefecture and its corresponding sample number

**Additional File 2.xls Table S2**. Summary of tick collection per species, life stage, and locations

**Additional File 3.xls Table S3**. Pairwise comparison of genetic differentiation (*F*_ST_) calculated for all *I. ovatus* using the *cox1* gene

**Additional File 4.xls Table S4.** Pairwise comparison of genetic differentiation (*F*_ST_) calculated for all *I. ovatus* using the 16S gene

**Additional File 5.xls Table S5.** Pairwise comparison of genetic differentiation (*F*_ST_) calculated for all *H. flava* using the *cox1* gene

**Additional File 6.xls Table S6**. Pairwise comparison of genetic differentiation (*F*_ST_) calculated for all *H. flava* using the 16S gene

**Additional File 7.docx Figure S1.** Plots a to d show the relationship between the pairwise *F*_ST_ values and the geographic distances for *I. ovatus* and *H. flava* in both *cox1* and 16S markers from the Mantel test analysis.

**Additional File 8.docx Figure S2.** An unweighted pair group method with the arithmetic mean (UPGMA) dendrogram of *I. ovatus* based on the pairwise genetic distance (*F*_ST_) of 16S among the 7 populations across Niigata Prefecture, Japan.

**Additional File 9.docx Figure S3** An unweighted pair group method with the arithmetic mean (UPGMA) dendrogram of *H. flava* based on the pairwise genetic distance (*F*_ST_) of *cox1* among the 5 populations across Niigata Prefecture, Japan.

**Additional File 10.docx Figure S4** An unweighted pair group method with the arithmetic mean (UPGMA) dendrogram of *H. flava* based on the pairwise genetic distance (*F*_ST_) of 16S among the 5 populations across Niigata Prefecture, Japan.

**Additional File 11.docx Figure S5** Median-joining haplotype network of *I. ovatus* from *cox1* marker. Each circle represents a unique haplotype and the lines correspond to mutations.

**Additional File 12.docx Figure S6** Phylogenetic tree from Bayesian analysis of the *cox1 I. ovatus* haplotype sequences with reference sequences from China

**Additional File 13.docx Figure S7** Median-joining haplotype network of *I. ovatus* from 16S marker. Each circle represents a unique haplotype and the lines correspond to mutations.

**Additional File 14.docx Figure S8** Median-joining haplotype network of *H. flava* from *cox1* marker. Each circle represents a unique haplotype and the lines correspond to mutations.

**Additional File 15.docx Figure S9** Median-joining haplotype network of *H. flava* from 16S marker. Each circle represents a unique haplotype, and the lines correspond to mutations.

**Additional File 16.docx Figure S10** Phylogenetic tree of 16S haplotype sequences of *I. ovatus* inferred by maximum likelihood analysis with evolutionary model GTR with sequences from China (MH208506, MH208512, MH208514 – MH208519, MH208522, MH208524, MH208531, MH208574, MH208577, MH208579, KU664519), Japan (AB819241, AB819242, AB819243, AB819244, and U95900). An outgroup (*Ixodes canisuga*) was also included.

**Additional File 17.docx Figure S11** Phylogenetic tree of *cox1* haplotype sequences of *H. flava* inferred by maximum likelihood analysis with evolutionary model HKY with sequences from China (KY021800-KY021807; KY021810-KY021819, KY003181, JQ62588 - JQ625889; JQ737097; JF758632). An outgroup (*Ixodes canisuga)* was also included.

**Additional File 18.docx Figure S12** Phylogenetic tree of 16S haplotype sequences of *H. flava* inferred by maximum likelihood analysis with evolutionary model GTR with sequences from China (KC844858 - KC844867; KC844880-KC844882, M696720, KP324926, KX450279) and Japan (AB19177-AB819192). An outgroup (*Ixodes canisuga)* was also included.

